# The Fragile X Messenger Ribonucleoprotein 1 regulates the morphology and maturation of human and rat oligodendrocytes

**DOI:** 10.1101/2024.08.16.608069

**Authors:** Vidya Ramesh, Ioanna Kougianou, Eleni Tsoukala, Zrinko Kozic, Karen Burr, Biju Viswanath, David Hampton, David Story, Bharath Kumar Reddy, Rakhi Pal, Owen Dando, Peter C. Kind, Sumantra Chattarji, Bhuvaneish T Selvaraj, Siddharthan Chandran, Lida Zoupi

## Abstract

The Fragile X Messenger Ribonucleoprotein (FMRP) is an RNA binding protein that regulates the translation of multiple mRNAs and is expressed by neurons and glia in the mammalian brain. Loss of FMRP leads to Fragile X Syndrome (FXS), a common inherited form of intellectual disability and autism. While most research has been focusing on the neuronal contribution to FXS pathophysiology, the role of glia, particularly oligodendrocytes, is largely unknown. FXS individuals are characterised by white matter changes which imply impairments in oligodendrocyte differentiation and myelination. We hypothesized that FMRP regulates oligodendrocyte maturation and myelination during postnatal development. Using a combination of human pluripotent stem cell - derived oligodendrocytes and an *Fmr1* knockout rat model, we studied the role of FMRP on mammalian oligodendrocyte development. We found that the loss of FMRP leads to shared defects in oligodendrocyte morphology in both rat and human systems *in vitro* which persist in the presence of FMRP expressing axons in chimeric engraftment models. Our findings point to species-conserved, cell-autonomous defects during oligodendrocyte maturation in FXS.

## Introduction

Oligodendrocytes are central nervous system glia that ensheath axons with myelin. Myelin is a lipid rich membrane that insulates and segregates the axons into functional domains enabling fast and energy-efficient signal transmission (1–3). Each oligodendrocyte targets and myelinates multiple axonal segments which belong to different neuronal subtypes and generates myelin sheaths that vary in length and in thickness (4–8). As such, the formation of each myelin sheath needs to be both spatially and temporally regulated to be able to adapt to the network’s requirements. This spatial and temporal resolution is achieved through local mRNA translation that depends on the function of multiple RNA binding proteins (9–13). Recent studies show that hundreds of mRNAs are localised in the myelin sheaths with *Fmr1* being identified as one of the highly enriched mRNAs in the nascent myelin sheaths in zebrafish (12, 14). Mutations in the *FMR1* gene lead to one of the most common single-gene causes of intellectual disability and autism, Fragile X Syndrome (FXS) (15).

FXS individuals experience a variety of symptoms including restrictive and repetitive behaviours, anxiety, hyperactivity, social avoidance, language impairments and high incidence of seizures (16). FXS is caused by a CGG repeat expansion in the 5’UTR region of the *FMR1* gene, leading to transcriptional silencing and loss of the fragile X messenger ribonucleoprotein (FMRP) from both neurons and glia in the brain (17, 18). FMRP is an RNA binding protein with multiple mRNA targets acting primarily as a translational repressor (17, 19–21). As such, a common feature among FXS preclinical models and human FXS-derived cells is the dysregulation of protein homeostasis that contributes to the development of several neuronal abnormalities (17, 22–26). Despite the important advances behind the molecular and cellular mechanisms underlying neuronal dysfunction, proposed neuron-focused treatments have not produced robust results in large clinical trials so far (27–31), while the contribution of glia in FXS pathophysiology remains largely understudied. FMRP is expressed throughout the oligodendrocyte lineage including mature oligodendrocytes in humans and in rodents (32–36). Furthermore, longitudinal imaging studies including in infants and in young children with FXS have identified alterations in white matter structures and changes in fibre densities when compared to typically developed individuals (37–39). Studies in *Fmr1* knockout mice showed significant delays during postnatal myelination in the cerebellum and hypomyelination of the auditory brainstem in the adult (40, 41). Furthermore, recent evidence in zebrafish showed that *Fmr1* is required for oligodendrocyte differentiation during larval stages and formation of myelin sheaths of appropriate length (14, 42). Despite the emerging evidence, it is yet unclear how FMRP regulates mammalian oligodendrocyte function.

We hypothesize that FMRP regulates oligodendrocyte maturation and myelination during postnatal development. To test our hypothesis, we used a combination of human pluripotent stem cell (hPSC) - derived oligodendrocytes (*FMR1* knockout and FXS individual-derived) and *Fmr1* knockout rats to assess the impact of FMRP loss on mammalian oligodendrocyte development. We found that the loss of FMRP leads to cell autonomous defects in oligodendrocyte maturation and morphology in both rat and human systems *in vitro*. Furthermore, using *ex vivo* and *in vivo* chimeric engraftment experiments, we show that these cell autonomous defects significantly impair their myelination potential *ex vivo* and morphology *in vivo*. Collectively our data identify oligodendrocyte specific dysregulations due to the loss of FMRP in both human and rat oligodendrocytes providing evidence that is of potential translational relevance to human FXS.

## Methods

### Animals

Rats and mice were housed and used in accordance with the guidelines established by the Animal Care (Scientific Procedures) Act 1986 and under the authority of Home Office Licenses in the UK and in accordance with the guidelines of the CPCSEA, Government of India and approved by the Institutional Animal Ethics Committees of the National Centre for Biological Sciences and the Institute for Stem Cell and Regenerative Medicine in India. The rats were male Long-Evans Hooded wild type and *Fmr1*^*em1/PWC*^ rats, hereafter referred to as *Fmr1*^*+/y*^ and *Fmr1*^*-/y*^ respectively (43). For the *in vitro* and *in vivo* analysis *Fmr1*^*+/y*^ and *Fmr1*^*-/y*^ rats were used. *Shiverer* mice (*Mbp*^*shi/shi*^ –C3Fe.SWV-*Mbp*^*Shi*^/J;001428) and *Rag1* mice (NOD.129S7(B6)-*Rag1*^*tm1Mom*^/J) were purchased from Jackson laboratories. *Mbp*^*Shi/+*^;*Rag1*^*-/-*^ males were bred with *Mbp*^*Shi/+*^ ;*Rag1*^*-/-*^ females to obtain *Mbp*^*shi/shi*^;*Rag1*^*-/-*^ male/female pups used for human OPC transplantation experiments. *Mbp*^*shi/shi*^ mouse pups of both sexes were used for the organotypic slice culture experiments. Both mice and rats were bred in-house and kept on a 12h/12h light/dark cycle with *ad libitum* access to food and water. Following weaning on postnatal day 21 (P21), rats and mice were group housed in mixed-genotype cages of 2-5 animals per cage. Animals transplanted with the same genotype of human oligodendrocyte precursor cells (hOPCs) were housed together.

### Rat oligodendrocyte progenitor cell (OPC) isolation

Cortices from neonatal rat pups (P0-P3) were isolated following dissection. The cortical tissue was digested using a papain solution, containing MEM (32360026, Life Technologies), N-Acetyl-L-cysteine (24mg/mL, A8199, Sigma-Aldrich), DNase type IV (40µg/mL, D5025, Sigma-Aldrich), papain (Worthington Biochemical 30U/mL) and incubated for 1-1.5 hours at 37°C and 5%CO_2_. Digested cortices were then washed with OPC growth medium containing DMEM (41966029, Life Technologies), 10% Fetal Bovine Serum (10270106, Life Technologies), 1% penicillin-streptomycin solution (15140122, Life Technologies) and triturated to single cell suspensions. Cell suspensions were transferred into Poly-D-lysine (1µg/mL, P6407, Sigma-Aldrich) coated 75cm^2^ flasks. The mixed glia cultures were kept in growth medium with frequent renewal every 48 hours for 10 to 14 days at 37°C and 5% CO_2_. OPC isolation was performed after sequential shaking for 1 hour at 220rpm and at 37°C for microglial removal, followed by further 16 hours at 37°C. The next day the OPC-enriched supernatants were incubated for further 25 minutes in 10cm petri dishes at 37°C and pooled according to genotype. 70,000 OPCs were seeded in 18mm, Poly-D-lysine coated glass coverslips in proliferation medium containing: DMEM (high glucose) + 0.5% Fetal Bovine Serum + 1% penicillin-streptomycin solution, 1% 100x SATO, 1% ITS liquid media supplement (I3146, Sigma-Aldrich), with freshly added PDGF-AA (10ng/ml, 100-13A, PeproTech) and FGF (10ng/ml, 100-18B, PeproTech) growth factors until day two *in vitro*. SATO medium contained 10 mg/mL BSA fraction V (A-4919, Sigma-Aldrich), 6µg/mL Progesterone (P-8783, Sigma-Aldrich), 1.61mg/mL Putrescine (P-5780, Sigma-Aldrich), 40µg/mL L-Thyroxine T4 (T-1775, Sigma-Aldrich) and 40µg/mL Tri-iodothyroxine (T-6397, Sigma-Aldrich). After two days, the proliferation growth factors were replaced by T3 (0.4µg/ml) and T4 (0.4µg/ml) to promote cell differentiation until day six *in vitro*.

### *Ex vivo Mbp*^*shi/shi*^ mouse cortical organotypic slices

P0–P3 *Mbp*^*shi/shi*^ mouse pups were decapitated and their brains were dissected in cold Hibernate™-A medium (A1247501, Thermo Scientific) on ice and the meninges were removed. The brains were immediately mounted on the vibratome and 250-300µm coronal cortical slices were collected in ice cold Hibernate™-A medium. The slices were immediately transferred onto Millicell cell culture inserts (30mm, hydrophilic PTFE, 0.4µm, PICM0RG50, Merck-Millipore) using a bent spatula and in warm slice medium containing 50% MEM (32360026, Life Technologies) with 25% Earle’s Balanced Salt Solution (24010043, Life Technologies), 25% heat-inactivated horse serum (26050088, Thermo Scientific), 1% Glutamax™ supplement (35050061, Thermo Scientific), 1% penicillin–streptomycin, 0.5% Amphotericin B (5290018, Thermo Scientific) and 6.5mg/ml glucose (G8769, Sigma-Aldrich). The slices were kept in serum-containing medium for approximately 5 days and gradually switched to serum-free medium containing DMEM/F12 (11320033, Thermo Scientific), 1% B-27™ supplement (17504044, Thermo Scientific), 0.5% N2 supplement (17502048, Thermo Scientific), 1% Glutamax™ supplement, 1% penicillin–streptomycin and 0.5% Amphotericin B before the addition of rat OPCs (44). Slices were kept at 37°C and 5% CO_2_ throughout the experiment. The medium was changed every two days. After approximately 10 days in culture 100,000 rat *Fmr1*^*+/y*^ or *Fmr1*^*-/y*^ cells were seeded on each cortical slice and cultured for two further weeks before fixation.

### Generation of human glial spheres and oligodendrocytes

The human embryonic stem cell line (male, Shef4, Supplementary Table 1), referred to as *FMR1*^*+/y*^ was obtained from the UK Stem Cell Bank (45). Gene-editing was performed on this line to generate the Shef4 *FMR1* knock-out hESC line referred to as *FMR1*^*-/y*^, as described previously (46). GM07072 (fragile X syndrome male, Supplementary Table 1) fibroblasts were obtained from the Coriell Institute for Medical Research under their consent and privacy guidelines as described (http://catalog.coriell.org). Induced human pluripotent stem cells were generated from this line, referred to as mutant FXS or mFXS, at Cedar-Sinai Medical Center (Los Angeles, CA) using standard protocols as previously described (47, 48). The mFXS line carrying the CGG repeat expansion was gene-corrected using previously published protocols (49, 50) to generate an isogenic control hiPSC line referred to as IsoFXS (Supplementary Table 1). All experiments were performed after obtaining statutory institutional ethical clearances. The characterization and validation of all human pluripotent stem cell lines used in this study were performed as described previously (47, 48, 51). Generation of oligodendrocyte cultures from hPSCs was performed according to previous published protocols (52). Embryo bodies were generated by cell lifting with Dispase 1mg/ml (17105-041, Life Technologies) and Collagenase 2mg/ml (17104-019, Life Technologies) and cultured for 7 days, with dual-SMAD inhibition, and in chemically defined media (CDM) that contained 50% Iscove’s modified Dulbecco’s medium (12440053, Thermofisher), 50% F12 medium (11765054, Thermofisher), BSA (5mg/ml, 05470, Sigma), 1% chemically defined Lipid concentrate (11905031, Gibco), monothioglycerol (450µM, M6145 Sigma-Aldrich), human insulin (7mg/ml, 11376497001, Roche), transferrin (15mg/ml, 10652202001, Roche), 1% penicillin/streptomycin (15140122, Gibco), supplemented with N-acetyl-L-cysteine (1mM, A7250, Sigma), Activin inhibitor SB43152 (10µM, 1614, Tocris), and LDN193189 (2µM, SML0559, Merck Millipore). Generated neurospheres were then caudalized by the addition of retinoic acid (1µM, R2625, Sigma) for a further 7 days. Spheres were ventralized with the addition of the sonic hedgehog agonist purmorphamine (1µM, 483367-10-8, Calbiochem) for 7 days in Advanced DMEM/F12 (Invitrogen) containing: 1% N-2 supplement (17502048, Invitrogen), 1% B27™ supplement (17504044, Invitrogen), 1% penicillin/streptomycin, 0.5% Glutamax™ supplement (35050061, Invitrogen), and 5µg/ml Heparin (Sigma). Spheres were further expanded in the presence of basic fibroblast growth factor (FGF)-2 (10ng/ml, 100-18B, PeproTech) for 7 days, following which neural differentiation was induced by FGF2 withdrawal for a further 2 weeks. The resultant glial spheres were further expanded for 2-4 weeks in oligodendrocyte proliferation media containing FGF2 (10ng/ml), PDGF-AA (20ng/ml, 100-13A, PeproTech), purmorphamine (1µM), SAG (1µM, 364590-63-6, Calbiochem), IGF-1 (10ng/ml, 100-11, PeproTech), T3 (60ng/ml, Sigma), and 1×ITS (41400045, Gibco) before initiating oligodendrocyte differentiation. Terminal differentiation of oligodendrocytes was achieved in oligodendrocyte differentiation media by mitogen withdrawal for 7 days (except for T3 and IGF-1) following single-cell dissociation using Papain (20 units/ml) dissociation kit (Worthington Biochemical) and plating on Matrigel (SLS, 354230; 1 in 100 dilution), fibronectin (20µg/ml, F2006, Sigma-Aldrich), laminin (10µg/ml, L2020, Sigma-Aldrich) coated coverslips at a density of 20,000-30,000 cells per 0.3cm^2^ until fixation.

To analyse the numbers of OPCs and proliferating cells in the glial spheres, 7-9-week-old spheres were plated on coverslips coated with Matrigel, Laminin and Fibronectin as mentioned above. 3-4 spheres were plated per coverslip and cultured for 2 days until fixation.

### Human OPC transplantation in *Mbp*^*shi/shi*^ mice pups

*Mbp*^*shi/shi*^;*Rag1*^*-/-*^ homozygous P0-P1 pups were anaesthetized with isoflurane and maintained on a heat mat during transplantation and during recovery. *FMR1*^*+/y*^ or *FMR1*^*-/y*^ glial spheres were dissociated at 7-9 weeks of age and injected into rostral and caudal positions within each cerebral hemisphere (70,000 cells per µl, 4 injections in total per animal) using a Hamilton^®^ syringe (24530, Sigma) and 30G needle. Injections were directed towards the corpus callosum and surrounding cortical grey matter. All pups in one litter were injected with the same genotype of OPCs. At 11-12 weeks of age, only the double homozygous animals were perfused under terminal anaesthesia using first ice-cold PBS, followed by fixation with 4% PFA (PFA, 252549, Sigma-Aldrich) and brains collected for histology.

### cDNA synthesis and qRT-PCR

RNA was extracted from 7–9-week-old glial spheres (*FMR1*^*+/y*^ and *FMR1*^*-/y*^) using the RNeasy micro kit from Qiagen (74004). RNA was converted to cDNA using RevertAid First Strand cDNA Synthesis Kit (K1621 Thermofisher). qPCR was performed using the DyNAmo ColorFlash SYBR Green qPCR Kit (F416 Thermofisher), CFX96 Touch Real-time PCR detection system (Biorad) and primers for human *FMR1* (Forward -TCCAATGGCGCTTTCTACA and Reverse -CATCATGAAATGGAATCTGCCTATC) and human *GAPDH* (Forward -TCGGAGTCAACGGATTTGGT and Reverse -TCCCGTTCTCAGCCTTGAC). Fold change was calculated using the Delta Delta Ct method (53).

### Immunofluorescence

#### Rat

Rats were perfused with 4% PFA and the brain tissue was harvested and post-fixed further with 4% PFA overnight at 4°C before transferred to 1xPBS solution. The frontal brain was sectioned using a Leica VT1000S. 100µm thick, coronal vibratome sections were briefly washed in PBS before blocking with 10% normal horse serum (26050088, Thermo Scientific), 0.3% Triton-X (X100, Sigma Aldrich) in 1xPBS for 2 hours at room temperature. For oligodendrocyte myelin sheath tracing experiments the sections were incubated in antigen unmasking solution (H-3300-250, Vector Laboratories) at 95°C for 20 minutes prior to blocking. Primary antibodies were diluted in the same blocking solution and sections were incubated for 48 hours at 4°C. After the primary antibody incubation, sections were washed in 1xPBS (3×1hr each), incubated with Alexa Fluor secondary antibodies (Thermo Fischer Scientific, 1:1000) for further 16 hours and counterstained with Hoechst 33342 solution (62249, Thermo Scientific) for nuclear visualization. All slides were mounted using Fluoromount-G^®^ mounting medium (0100-01, Southern-Biotech).

For immunocytochemistry, the glass coverslips were fixed with 4% PFA for 20 minutes at room temperature and then washed three times with 1xPBS. Following permeabilization for 10 minutes at room temperature and in 0.1% Triton-X in 1xPBS, coverslips were blocked for 30 minutes in 10% normal horse serum in 1xPBS at room temperature. The primary antibodies were diluted in the same blocking solution and coverslips were incubated for 2 hours at room temperature. Subsequently, the coverslips were washed in 1xPBS, incubated with secondary antibodies for 1.5 hours at room temperature and counterstained with Hoechst 33342 solution before mounting on glass slides with Fluoromount-Gscript® mounting medium.

For organotypic slice immunohistochemistry the slices were washed once with 1xPBS before they were fixed with 4% PFA for 1 hour at room temperature. The slices were subsequently rinsed in 1xPBS and blocked in 3% heat-inactivated horse serum, 2% BSA (A7906, Sigma-Aldrich), and 0.5% Triton X-100 in 1xPBS for 2 hours at room temperature. Following blocking, the slices were incubated for 48 hours with primary antibodies diluted in blocking solution at 4°C. The slices were then washed 3 times with blocking solution and incubated with the appropriate secondary antibodies overnight at 4°C. Finally, the slices were washed in 1x PBS, counterstained with Hoechst 33342 solution and mounted on a glass microscopic slide using Fluoromount-G^®^ mounting medium.

The primary antibodies used were as follows: against MBP (rat monoclonal, MCA409S, BioRad, 1:300), Neurofilament-H (chicken polyclonal anti-NF-H, 822601, Biolegend. 1:10.000), OLIG2 (rabbit polyclonal, AB9610, Sigma-Aldrich, 1:100), CNPase (mouse monoclonal, AMAB91072, Sigma-Aldrich, 1:2000), PDGFRα (goat polyclonal, AF1062-SP, Novus Biologicals, 1:200), CC-1 (mouse monoclonal, OP80 | Anti-APC (Ab-7) Mouse mAb (CC-1), OP80, EMD Millipore 1:300).

### Human

For transplant experiments, mice were perfused with 4% PFA, and the brain was post-fixed with 4% PFA overnight at 4°C before being transferred to 30% sucrose/PBS solution for cryoprotection. Brains were frozen in OCT freezing media (Leica, 14020108926) and sectioned using a Leica CM1850 cryostat. Sagittal sections of 16µm thickness were obtained and kept frozen until use. For immunohistochemistry, slides were briefly washed in 1xPBS before permeabilizing with 0.5% Triton-X in 1xPBS for 15 minutes followed by blocking with 10% normal goat serum (S1000, Vector labs), 0.1% Triton-X in 1xPBS for 1 hour at room temperature. Primary antibodies were diluted in the same blocking solution and sections were incubated for 16-20 hours at 4°C. After the primary antibody incubation, sections were washed in 1xPBS (3×15min each) and incubated with Alexa Fluor secondary antibodies (Thermo Fischer Scientific, 1:1000) for 1 hour at room temperature. Following 3×15min washes with 1xPBS, the slides counterstained with DAPI (D9542, Sigma) for nuclear visualization. All slides were mounted using Fluorsave mounting medium (345789, Merck Millipore) and 24×50mm glass coverslips.

For immunocytochemistry, 7-day old oligodendrocyte cultures or 2-day old glial spheres plated on glass coverslips were fixed with 4% PFA for 15 minutes at room temperature and then washed three times with 1xPBS. Following permeabilization for 10 minutes at room temperature with 0.2% Triton-X in 1xPBS, immunocytochemistry was performed in the same way as the transplant immunostaining. For PDGFRα and O4 staining, the primary antibodies were added on live cells for 2 hours prior to PFA fixation.

The primary antibodies used were as follows: SOX10 (Rabbit polyclonal, AB155279, Abcam, 1:1000), OLIG2 (Rabbit polyclonal, AB9610, Millipore, 1:1000), PDGFRα (Rabbit polyclonal, D13C6, Cell Signaling), MBP (Rat monoclonal, AB7349, Abcam, 1:50), O4 (MAB1326, R&D systems), Human Nuclei (Mouse monoclonal, AB1281, Millipore, 1:500), Ki67 (Rabbit polyclonal, AB9260, Merck Millipore, 1:500).

### Image acquisition and analysis

Imaging was done either using Leica TCS SP8, Leica Thunder Imager, Olympus FV1000 or Zeiss LSM710 confocal microscopes.

#### Rat analysis

For cell density quantification 3-6 z-stacks of 40µm thickness (63× objective, 184.58µm x 184.58µm each; pixel size 180.38nm x 180.38nm, step size: 0.5µm) were obtained from layers 2/3 of the medial prefrontal cortex from both hemispheres. Oligodendrocyte lineage cell numbers and total cell numbers were quantified automatically using IMARIS 3D imaging software (RRID:SCR_007370, spots module) and represented as % of cells/total population. From each rat 3 sections were analysed on average and the numbers were averaged for each animal.

The average sheath number and myelin sheath length were quantified as previously shown (54). Briefly, using the same microscope settings as above, we imaged individual CNPase positive oligodendrocytes of the layer 2/3 of the medial prefrontal cortex in the whole 100µm vibratome section (step size 0.5µm and 1024 x 1024 resolution). The intermodal length was quantified using the Simple Neurite Tracer plugin and the sheath number was subsequently quantified using the cell counter plug-in in Fiji image analysis software (Fiji, RRID:SCR_002285). The intermodal lengths were binned and a percentage of frequency distribution is depicted. Alternatively, the log_10_ of the lengths was calculated and represented as a % percent frequency.

For the *in vitro* analysis 3-4 coverslips were imaged per genotype and per experiment. Each coverslip was scanned using 63×objective and 1024×1024 resolution. Composite areas consisting of 45-65 tiles of 184.52µm x 184.52µm each were obtained for the cell density quantification *in vitro*. Cell densities were quantified with Fiji image software. For the Sholl analysis z-stacks of individual oligodendrocytes (20-40 cells/genotype/experiment and 4 different experiments) were imaged and analysed using the Sholl analysis plugin in Fiji with 10µm concentric circle intervals. The total number of intersections relative to the distance of the cell body is shown.

#### Human analysis

For the transplant analysis, sections were imaged using 40x objective and 1024×1024 resolution (16µm-thick sections, step size 0.5µm). Areas containing human nuclei in the corpus callosum, and fornix were imaged with 2-3 FOV per section. From each mouse, 2-3 sections were analysed, and the numbers were averaged for each animal. Analysis was performed using a CellProfiler (RRID:SCR_007358) pipeline wherein the MBP image threshold was set and the MBP area was quantified per MBP cluster. 10-20 MBP clusters were analysed per animal. For SOX10 analysis, a CellProfiler automated analysis pipeline was used to quantify colocalised SOX10 and human nuclei and total human nuclei per FOV. For the *in vitro* Sholl analysis 2-3 coverslips were imaged per genotype and per experiment. Each coverslip was scanned using 40x objective, 1.5 zoom and 1024×1024 resolution (step size 0.5µm). For the Sholl analysis z-stacks of individual oligodendrocytes (5-15 cells/genotype/experiment and 4 different experiment) were imaged and analysed using the Sholl analysis plugin in Fiji software with 2µm concentric circle intervals. The total number of intersections relative to the distance of the cell body is shown.

For % OLIG2, Ki67, PDGFRα and SOX10 over total nuclei analysis in the glial spheres, only cells which had migrated out of the spheres after 2DIV were analysed. 2 coverslips were imaged per genotype and per experiment. Each coverslip was scanned using 20x objective (step size 1µm, 1024×1024 resolution). The marker positive cells and total nuclei (DAPI positive) were counted manually using the cell counter plugin in Fiji. For %SOX10/total cells analysis in 7-day old oligodendrocyte cultures, a CellProfiler automated analysis pipeline was used to identify SOX10 and DAPI nuclei.

### *In silico* analysis

We downloaded the single cell sequencing dataset of oligodendrocyte lineage cells from (35). Expression data were imported into Seurat (55) (R package version 4.3.0), and the thirteen distinct populations of cells in the data set were consolidated into six major classes: OPC (oligodendrocyte precursor cells), COP (differentiation-committed oligodendrocyte precursors), NFOL (newly formed oligodendrocytes), MFOL (myelin-forming oligodendrocytes), MOL (mature oligodendrocytes), and VLMC (vascular and leptomeningeal cells); VLMCs were excluded from further analysis. Per-cell expression values were normalized using Seurat’s “LogNormalize” method, and average gene expression calculated per-cell class. A set of 7347 expressed genes was constructed, after filtering out genes expressed lower than an average of 1 in 100,000 counts per-cell.

We then intersected these genes with HITS-CLIP data of FMRP targets in the postnatal (P11-25) mouse brain (56) to form a background universe of 7066 genes which were both expressed in the single-cell data and analysed by Maurin et al., and for each oligodendrocyte cell class, ranked genes by their average expression in that class. We performed gene ontology (GO) enrichment analysis of the 2285 FMRP targets (those genes with PEAKS > 0 as identified in Maurin et al.) in our gene universe, in the biological process category, using topGO (57) (R package version 2.52.0), retaining those GO terms enriched with p-value < 0.01, and with at least ten FMRP target genes. For each such GO term, we performed a Kolmogorov-Smirnov test on the ranks of the FMRP targets for that term, in the ranked expression list for each cell class, to determine skewedness of ranks (where the more positive the KS-test statistic, the more skewed the ranks of the FMRP targets are towards the top of the expression ranking list for a cell class).

### Statistical analysis

Statistical analysis was performed using GraphPad Prism 9 or 10 (RRID:SCR_002798). Data were tested for normal distribution using the D’Agostino–Pearson test or the Shapiro–Wilk test. The variance of the data was assessed using the F test for variance. Depending on data distribution parametric or nonparametric tests were used. A difference was considered statistically significant when p<0.05. Data are shown as mean ± sem or mean ± sd. Details of statistical test used, precise p values and n values for each comparison are detailed in the main text and figure legends. Illustrations created with BioRender.com.

## Results

### Impaired morphology and decreased maturation of rat *Fmr1*^*-/y*^ oligodendrocytes *in vitro*

The observed white matter defects in FXS children and in animal models point to dysregulations in myelination and oligodendrocyte function early in postnatal development. As FMRP is expressed by oligodendrocytes (33, 35, 36), we first sought to identify its potential targets within the oligodendrocyte lineage. We integrated a single-cell sequencing dataset of oligodendrocyte lineage cells (35) with HITS-CLIP data of FMRP targets from the postnatal mouse brain (56). FMRP targets were expressed throughout the oligodendrocyte lineage. To explore the oligodendrocyte-specific processes that are likely to be disrupted by the loss of FMRP, we subsequently performed a GO enrichment analysis on these targets (Supplementary Table 2). Enriched GO terms included processes involved in synaptic assembly, cytoskeletal and microtubule organisation, glucose homeostasis and regulation of myelination that are known to be implicated in different stages of oligodendrogenesis and differentiation (58–68). For most GO terms, oligodendrocyte progenitor cells (OPCs) showed the highest enrichment in potential mRNA binding targets within the oligodendrocyte lineage (Supplementary Figure 1).

We then sought to determine the oligodendrocyte-specific effects of FMRP loss at the cellular level in mammalian systems. We isolated OPCs from the cortices of *Fmr1*^*+/y*^ and *Fmr1*^*-/y*^ rat pups and cultured them in the absence of neurons for six days *in vitro* (Figure 1A,B,E). Following immunocytochemistry for transcription factor OLIG2 and for myelin basic protein (MBP) (Figure 1A,B), we first quantified the percentage of OLIG2 expressing oligodendrocyte lineage cells in our cultures and found no difference between genotypes (*Fmr1*^*+/y*^: 70.06% ± 4.926 OLIG2+ cells/total cells, n=5 experiments, *Fmr1*^*-/y*^: 67.82% ± 5.237 OLIG2+ cells/total cells, n=3 experiments; p= 0.5806) (Figure 1F). In contrast, when we measured the percentage of mature oligodendrocytes (OLIG2+MBP+) we found a significant decrease in the *Fmr1*^*-/y*^ cultures compared to controls (*Fmr1*^*+/y*^: 70.70% ± 8.392 OLIG2+MBP+ cells/OLIG2+ cells, n=5 experiments, *Fmr1*^*-/y*^: 36.33% ± 9.051 OLIG2+MBP+ cells/OLIG2+ cells, n=3 experiments; p= 0.0016) which suggests an effect during oligodendrocyte maturation and not during the generation of oligodendrocyte lineage cells *in vitro*. We also observed that the morphology of the *Fmr1*^*-/y*^ MBP-expressing oligodendrocytes was different to those of the *Fmr1*^*+/y*^ oli-godendrocytes (Figure 1C, D). To assess the morphological features of mature oligodendrocytes in our cultures we performed Sholl analysis on individual MBP-expressing oligodendrocytes after six days in culture (Figure 1H). *Fmr1*^*-/y*^ oligodendrocytes showed simpler morphologies and fewer branching points *in vitro* than the wild type oligodendrocytes (Genotype effect: F = 25.85. DFn = 1, DFd = 50, p < 0.0001, n=4 different experiments/genotype). These results point to an oligodendrocyte-specific defect in the late maturation states due to the loss of FMRP.

**Fig. 1.**
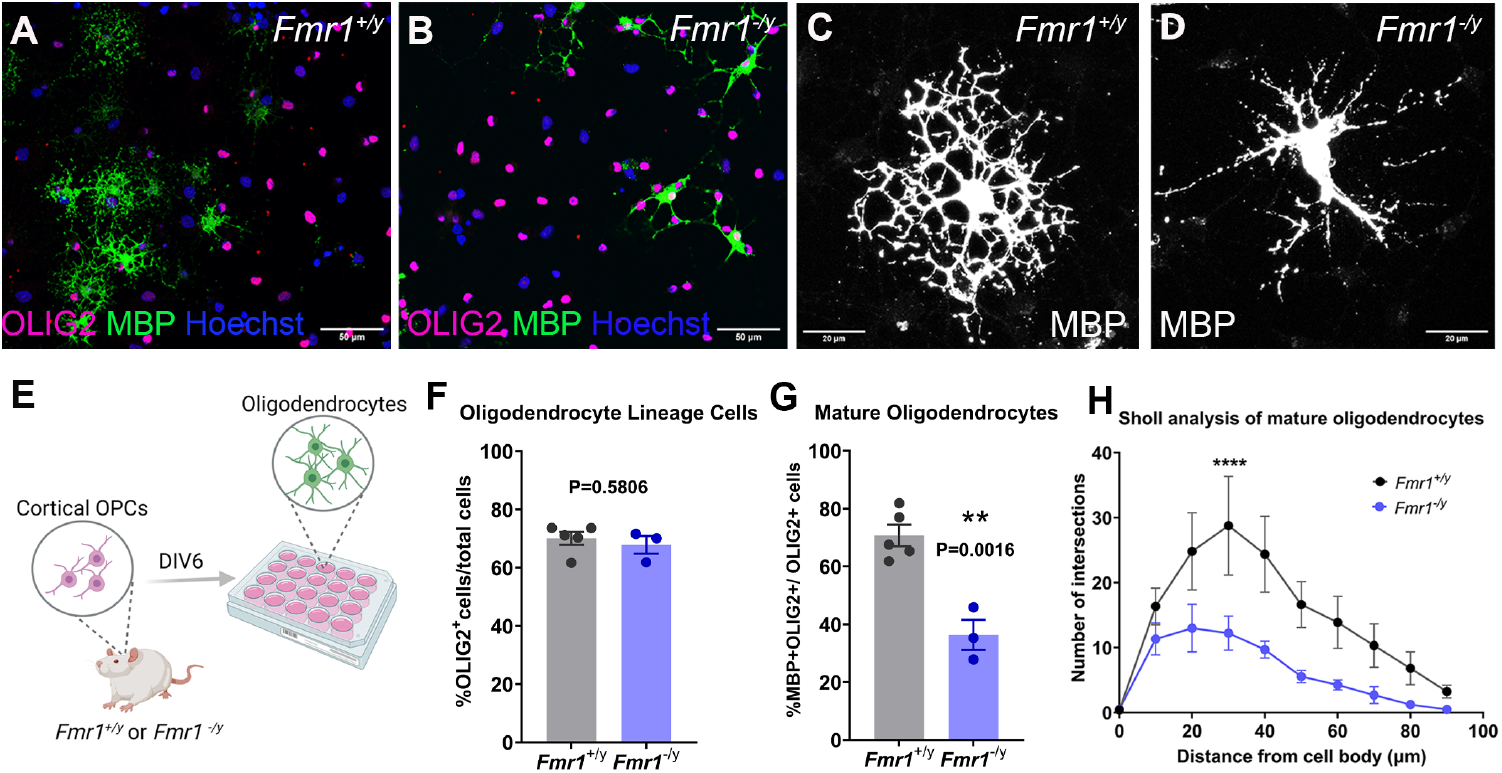
Impaired morphology and maturation of rat *Fmr1*^*-/y*^ oligodendrocytes *in vitro*. **A-B:** Representative images of cortical oligodendrocytes after 6 days *in vitro* immunostained for OLIG2 (magenta), MBP (green) and counterstained nuclei with Hoechst (blue). **C-D:** Representative images of MBP-expressing mature oligodendrocytes from each genotype after 6 days *in vitro*. **E:** Experimental design. **F:** Percent of OLIG2-expressing cells over the total cell number after 6 days *in vitro*; data presented as mean±sem and each circle is an independent experiment. **G:** Percent of MBP-expressing cells over the total OLIG2+ cell number after 6 days *in vitro*; data presented as mean±sem and each circle is an independent experiment. **H:** Sholl analysis of oligodendrocyte morphology between genotypes; data presented as mean±sem of 4 different experiments. P value calculated in F,G with two-tailed, unpaired t-test with Welch’s correction and with 2-Way ANOVA with Šídák’s multiple comparisons test in H. ^****^ indicates p<0.0001.

### Impaired morphology of human *FMR1*^*-/y*^ and mFXS oligodendrocytes *in vitro*

To determine if the cellular defects of rat *Fmr1*^*-/y*^ oligodendrocytes were also conserved in human oligodendrocytes that lacked FMRP, we generated human embryonic stem cell (hESCs) derived oligodendrocytes using a previously published protocol from our group (52). Glial precursor-enriched spheres were derived from *FMR1* null hESCs (named *FMR1*^*-/y*^) and from an isogenic hESC control (named *FMR1*^*+/y*^) and differentiated into glial cultures containing oligodendrocytes (Figure 2A). The downregulation of *FMR1* which has been previously described by our group (48), was confirmed by quantitative RT-PCR in glial spheres (Figure 2B). First, we investigated if there were any changes in the generation of oligodendrocyte precursors using immunocytochemistry for OLIG2 and the cell surface marker platelet-derived growth factor receptor alpha (PDGFRα) across genotypes and found no difference (Supplementary Figure 2A-G) (*FMR1*^*+/y*^: 51.62% ± 5.51 OLIG2+cells/total cells, and *FMR1*^*-/y*^: 49.69% ± 5.99 OLIG2+ cells/total cells, n=4 experiments, p=0.82; *FMR1*^*+/y*^: 13.3% ± 1.36 PDGFRα+ cells/total cells and *FMR1*^*-/y*^: 20.17% ± 7.29 PDGFRα+ cells/total cells, n=3 experiments, p=0.44). We further analysed cell proliferation using immunocytochemistry for Ki67, a nuclear proliferation marker, and found no difference across genotypes (Supplementary Figure 2H-J) (*FMR1*^*+/y*^: 37.34% ± 0.24 Ki67+ cells/total cells and *FMR1*^*-/y*^: 31.43% ± 1.45 Ki67+ cells/total cells, n=3 experiments, p=0.052). Following this, we generated glial cultures containing oligodendrocytes from *FMR1*^*-/y*^ hESCs and *FMR1*^*+/y*^ hESCs as well as FXS individual-derived hiPSCs (mFXS) and isogenic hiPSC control (IsoFXS) (Figure 2A and Supplementary table 1). This FXS-derived iPSC line has been previously characterised by our group to show downregulation of FMRP protein compared to its gene-edited control (47). Using immunocytochemistry for SOX10 in seven-day old glial cultures we tested for possible changes in the proportions of oligodendrocyte lineage cells and observed no difference across genotypes (Supplementary Figure 3) (*FMR1*^*+/y*^: 18.7% ± 9.41 SOX10+ cells/total cells and *FMR1*^*-/y*^: 16.5% ± 9.74 SOX10+ cells/total cells, n=3 experiments, p=0.88; mFXS 19% ± 9.96 SOX10+ cells/total cells and IsoFXS 35.3% ± 8.88 SOX10+ cells/total cells, n=3 experiments, p=0.29). Human oligodendrocytes at this stage (seven days in culture) express O4 which allowed us to assess their morphology using Sholl analysis (Figure 2C-F). *FMR1*^*-/y*^ and mFXS oligodendrocytes showed fewer branching points *in vitro* than the control *FMR1*^*+/y*^ and IsoFXS oligodendrocytes respectively (Figure 2G,H) (*FMR1*^*+/y*^ and *FMR1*^*-/y*^: Genotype effect: F = 27.07, DFn = 1, DFd =256, p < 0.0001, n=4 experiments/genotype; IsoFXS and mFXS: Genotype effect: F = 24.72, DFn = 1, DFd =283, p<0.0001, n=4 experiments/genotype) similar to the effect observed in rat *Fmr1*^*-/y*^ oligodendrocytes. These data suggest that there is a shared role for FMRP on oligodendrocyte maturation and morphology across species.

**Fig. 2.**
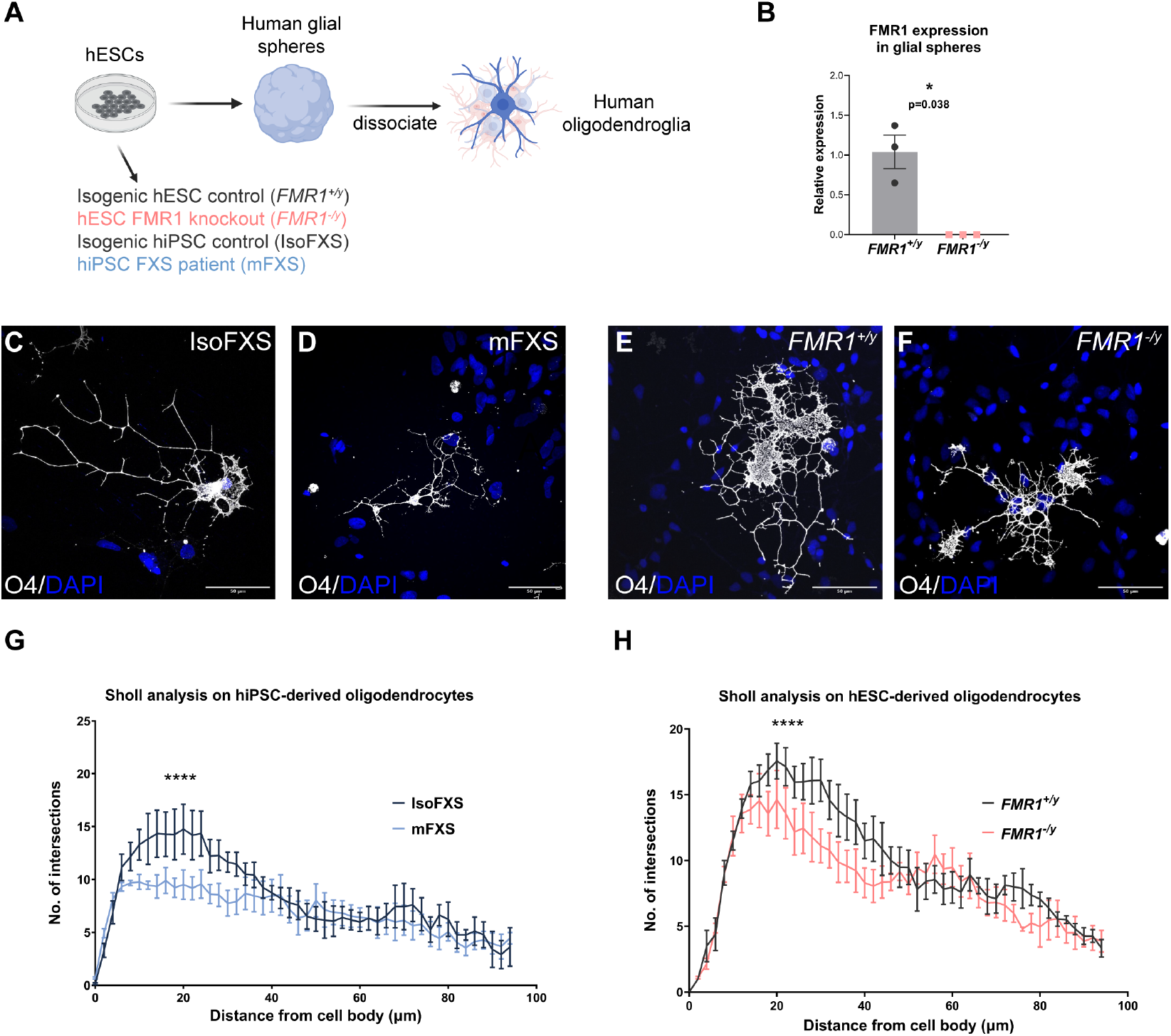
Impaired morphology of human *FMR1*^*-/y*^ and mFXS oligodendrocytes *in vitro*. **A:** Schematic showing generation of human mixed glial cultures containing oligodendrocytes (dark blue), OPCs/NPCs (light blue) and astrocytes (pink) **B:** qRT-PCR for *FMR1* mRNA in 7-9 week old *FMR1*^*+/y*^ and *FMR1*^*-/y*^ glial spheres; each data point is an independent experiment **C-F:** Representative images of *FMR1*^*+/y*^, *FMR1*^*-/y*^, IsoFXS and mFXS Human oligodendrocytes after 7 days *in vitro* immunostained for O4 (white) and counterstained nuclei with DAPI (blue) **G:** Sholl analysis of oligodendrocyte morphology between IsoFXS and mFXS oligodendrocytes. **H:** Sholl analysis of oligodendrocyte morphology between *FMR1*^*+/y*^ and *FMR1*^*-/y*^ oligodendrocytes. Data points and error bars represent means and sem respectively from 4 different experiments; P-value in G,H calculated using 2-way ANOVA and in B using two-tailed unpaired t-test with Welch’s correction. ^****^ indicates p<0.0001.

### Cortical myelination impairment in the prefrontal cortex of *Fmr1*^*-/y*^ rats *in vivo*

Our analysis of human and rat *Fmr1*-null oligodendrocytes identified morphological and maturation defects *in vitro*. To determine if these impairments are also evident *in vivo*, we examined the sparsely myelinated layers 2/3 of the medial prefrontal cortex (mPFC) in *Fmr1*^*+/y*^ and *Fmr1*^*-/y*^ rats in postnatal development when myelination was still ongoing. Dysfunction of the PFC, a region that is involved in emotional behaviour and cognitive functions, has been associated with cognitive impairments in both FXS individuals (69–72) and in *Fmr1*^*-/y*^ rodents (43, 73–75). Furthermore, changes in mPFC myelination have been linked to impaired memory and social interactions (76–79).

We first tested for changes in oligodendrocyte lineage cell densities during the third postnatal week (early mPFC myelination) using immunohistochemistry for PDGFRα (OPCs), APC (CC-1, mature oligodendrocytes) and OLIG2 (oligodendrocyte lineage cells) in layers 2/3 of the mPFC (Supplementary Figure 4). We did not observe any significant differences in the densities of oligodendrocyte progenitors (PDGFRα+ cells/OLIG2+ cells *Fmr1*^*+/y*^: 0.6868 ± 0.03370, n=5 rats, *Fmr1*^*-/y*^: 0.6952± 0.01823, n=4 rats, p=0.6494) (Supplementary Figure 4A), of oligodendrocyte lineage cells (OLIG2+ cells/total cells *Fmr1*^*+/y*^: 0.1311 ± 0.02748, n=5 rats, *Fmr1*^*-/y*^: 0.1421± 0.01342, n=4 rats, p=0.4629) (Supplementary Figure 4B), or of mature oligodendrocytes (CC1+ cells/OLIG2+ cells *Fmr1*^*+/y*^: 0.07847 ± 0.01690, n=5 rats, *Fmr1*^*-/y*^: 0.07072 ± 0.01656, n=4 rats, p=0.5124) (Supplementary Figure 4C) oligodendrocyte cell populations are present in the absence of FMRP at the early stages of mPFC myelination *in vivo* in the rat.

To assess changes during postnatal mPFC myelination *in vivo*, we next asked if the *Fmr1*^*-/y*^ myelin forming oligodendrocytes generate comparable numbers of myelin sheaths and of similar internodal lengths to the *Fmr1*^*+/y*^ oligodendrocytes. To address this question we used a previously established immunohistochemistry method (54) that allows the tracing of individual oligodendrocyte cell bodies, their processes and their myelin sheaths using a CNPase antibody (Figure 3A,B). We traced individual oligodendrocytes in layers 2/3 of the mPFC and assessed their morphologies in two different postnatal timepoints (3rd and 5th postnatal week). Our analysis showed that the *Fmr1*^*-/y*^ oligodendrocytes form on average significantly fewer myelin sheaths than the wild types (*Fmr1*^*+/y*^: 31.85 ± 2.016 average sheaths/oligodendrocyte, n=6 rats, *Fmr1*^*-/y*^: 23.41± 4.329 sheaths/oligodendrocyte, n=5 rats; p= 0.0087), but of comparable lengths (*Fmr1*^*+/y*^: 41.66 ± 5.429 µm,1200 sheaths, n=4 rats, *Fmr1*^*-/y*^: 40.64 ± 5.911 µm, 827 sheaths n=3 rats; p= 0.8274) during the third postnatal week (Figure 3C,D). However, no difference was detected between genotypes at the fifth postnatal week for both the mean myelin sheath number per oligodendrocyte (*Fmr1*^*+/y*^: 35.93 ± 4.597 sheaths/oligodendrocyte, n=8 rats, *Fmr1*^*-/y*^: 35.10 ± 5.339 sheaths/oligodendrocyte, n=8 rats; p=0.7450) and for the mean intermodal length (*Fmr1*^*+/y*^: 50.80 ± 5.313µm,1280 sheaths, n=3 rats, *Fmr1*^*-/y*^: 50.59 ± 3.231 µm, 1354 sheaths n=3 rats; p= 0.9569) (Figure 3E,F). These results show that in addition to the altered oligodendrocyte morphology seen *in vitro, Fmr1*^*-/y*^ oligodendrocytes show early defects in myelin sheath formation *in vivo* which are in line with previous findings showing early myelination deficits in the cerebellum of *Fmr1* null mice (40).

**Fig. 3.**
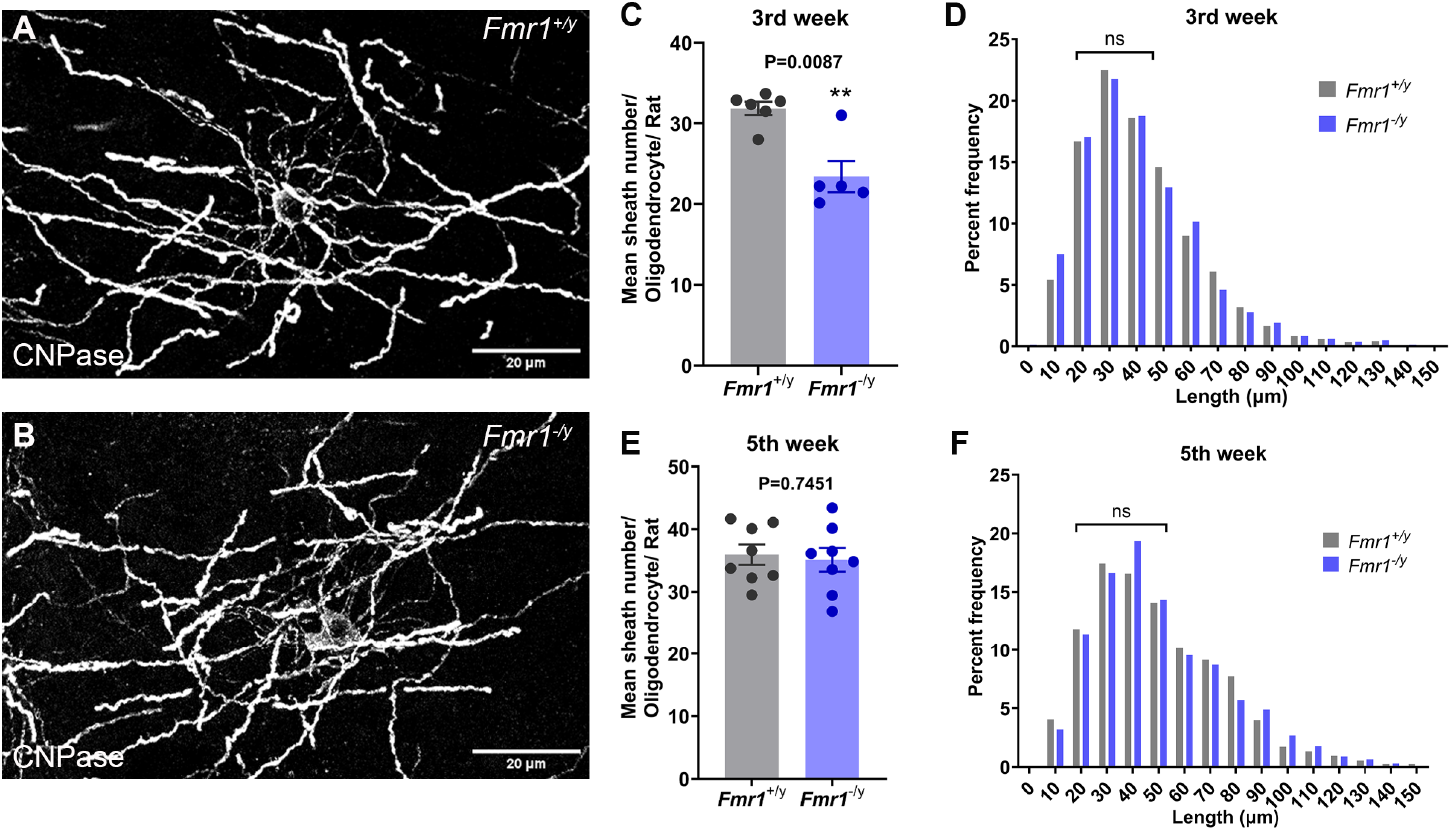
Cortical myelination impairment in the prefrontal cortex of *Fmr1*^*-/y*^ rats. **A-B:** Example of myelinating oligodendrocytes at the 3rd postnatal week in the rat mPFC stained for CNPase. **C:** Mean myelin sheath number/oligodendrocyte in layers 2/3 of the mPFC during the 3rd postnatal week. **D:** Percent frequency of myelin sheath length distribution per genotype at the 3rd postnatal week. **E:** Mean number/oligodendrocyte in layers 2/3 of the mPFC during the 5th postnatal week. **F:** Percent frequency of myelin sheath length distribution per genotype at the 5th postnatal week. Each data point is a rat. Data presented as mean±sem. P values calculated with two-tailed Mann Whitney test for C and two-tailed, unpaired t-test for the mean sheath number in E and mean sheath length in D and F.

### Impaired morphology of human *FMR1*^*-/y*^ oligodendrocytes *in vivo* in hypomyelinated immunosuppressed mice

To assess if the *in vitro* morphological defects observed in *FMR1*^*-/y*^ human oligodendrocytes are cell-autonomous and retained *in vivo*, we employed an *in vivo* chimeric model wherein dissociated *FMR1*^*+/y*^ or *FMR1*^*-/y*^ glial spheres were transplanted into neonatal immunodeficient and hypomyelinated MBP-deficient mice (*Mbp*^*shi/shi*^; *Rag1*^*-/-*^) that normally express FMRP. Since *Mbp*^*shi/shi*^ mice lack expression of MBP protein (80, 81) any MBP expression would derive from the transplanted human oligodendrocytes. P0-P1 neonatal pups were injected with *FMR1*^*+/y*^ or *FMR1*^*-/y*^ dissociated spheres containing OPCs (4 injections of 70,000 cells per injection) into the rostral and caudal neocortex and analysed at 11-12 weeks post transplantation (Figure 4A). Human nuclei were largely found in the areas surrounding the lateral ventricles (mainly in the corpus callosum) (Figure 4B,C) and in the fornix. To identify the human cells which differentiated into oligodendrocytes, we performed immunohistochemistry for MBP and traced the clusters of MBP transplanted cells *in vivo* (Figure 4D-F). Although at this stage, the differentiated human oligodendrocytes had not made distinct myelin sheaths (82) we were able to assess their MBP+ oligodendrocyte morphology. Analysis of the MBP+ area per cluster revealed a reduction in relative area in *FMR1*^*-/y*^ oligodendrocytes compared to *FMR1*^*+/y*^ oligodendrocytes (*FMR1*^*+/y*^ :6233 ± 652.6, *FMR1*^*-/y*^ :3520 ± 255.8, n=3 mice, p=0.039). In contrast, the numbers of human SOX10-positive oligodendrocyte lineage cells in the *Mbp*^*shi/shi*^; *Rag1*^*-/-*^ brains were comparable between genotypes (*Fmr1*^*+/y*^ : 31.36% ± 5.1784 SOX10+ cells/total Human nuclei+ cells, *Fmr1*^*-/y*^ : 22% ± 5.98 SOX10+ cells/total Human nuclei+ cells, n=3 mice, p=0.31), (Figure 4G-I) suggesting that the difference in MBP expression is likely due to impaired maturation and not due to a difference in oligodendrocyte lineage cell numbers. These data indicate that the *FMR1*^*-/y*^ human oligodendrocytes have reduced MBP expression among mouse axons that express FMRP.

**Fig. 4.**
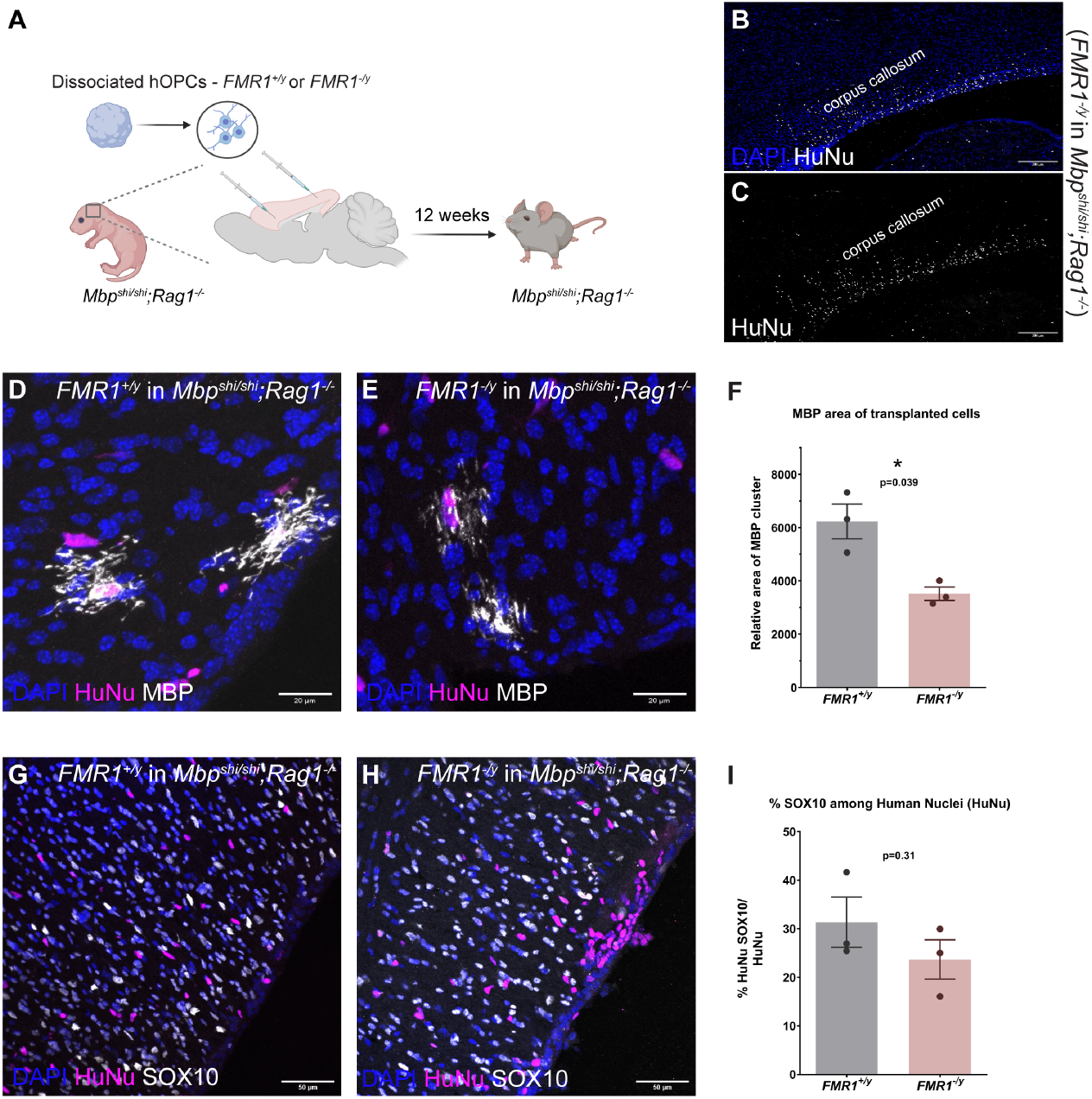
Impaired morphology of transplanted human *FMR1*^*-/y*^ oligodendrocytes *in vivo* in *Mbp*^*shi/shi*^,*Rag1*^*-/-*^ (immunosuppressed shiverer) mice. **A:** Schematic of chimeric transplantation paradigm **B-C:** Representative image showing spread of human cells in the corpus callosum of *Mbp*^*shi/shi*^,*Rag1*^*-/-*^ mice immunostained for Human Nuclei marker (HuNu, white) and counterstained with DAPI (blue). **D-E, G-H:** Representative images of transplanted *FMR1*^*+/y*^ and *FMR1*^*-/y*^ human oligodendrocytes in *Mbp*^*shi/shi*^,*Rag1*^*-/-*^ mice at 12 weeks immunostained for MBP (white), Human Nuclei (magenta), SOX10 (white), and counterstained with DAPI (blue) **F:** Graph showing relative MBP volume per MBP cluster of *FMR1*^*+/y*^ and *FMR1*^*-/y*^ transplanted oligodendrocytes. **I:** Graph showing % SOX10 among Human Nuclei population in *FMR1*^*+/y*^ and *FMR1*^*-/y*^ transplanted shiverer mice. Each data point is a mouse from a different litter. Error bars indicate sem from 3 different experiments; P values calculated using two-tailed unpaired t-test with Welch’s correction.

### Rat *Fmr1*^*-/y*^ oligodendrocytes form fewer myelin sheaths on *Mbp*^*shi/shi*^ axons *ex vivo*

Given that rat *Fmr1*^*-/y*^ oligodendrocytes exhibit morphological and maturation deficits *in vitro* and myelination impairments *in vivo*, we sought to determine if these deficits are cell-autonomous and retained in the presence of FMRP-expressing axons as in the case of human *FMR1*^*-/y*^ oligodendrocytes. To assess this, we used an *ex vivo* chimeric co-culture system in which *Fmr1*^*+/y*^ or *Fmr1*^*-/y*^ rat oligodendrocyte progenitors were transplanted onto cortical organotypic slices derived from neonatal *Mbp*^*shi/shi*^ mice. *Mbp*^*shi/shi*^ cortices were seeded with 100,000 oligodendrocyte progenitor cells and analysed after 2 weeks of co-culture to match the early stages of cortical myelination *in vivo* (Figure 5A). The seeded rat oligodendrocytes were detected both in the corpus callosum and in the cortex using immunohistochemistry for MBP (Figure 5B,C). Individually traced *Fmr1*^*-/y*^ rat oligodendrocytes on *Mbp*^*shi/shi*^ cortices formed significantly fewer myelin sheaths than *Fmr1*^*+/y*^ oligodendrocytes after two weeks in culture (*Fmr1*^*+/y*^: 27.19± 6.189 sheaths/oligodendrocyte, n=3 experiments, *Fmr1*^*-/y*^: 14.14± 3.293 sheaths/oligodendrocyte, n=3 experiments; p= 0.0473) (Figure 5D) but of comparable internodal lengths (*Fmr1*^*+/y*^: 42.84 µm ± 20.02, 1168 sheaths, n=4 experiments, *Fmr1*^*-/y*^: 42.77 µm ± 17.79, 1005 sheaths n=4 experiments; p= 0.9960) (Figure 5E,F). This indicates that the *Fmr1*^*-/y*^ rat oligodendrocytes form fewer myelin sheaths than wildtypes, on mouse axons that express FMRP.

**Fig. 5.**
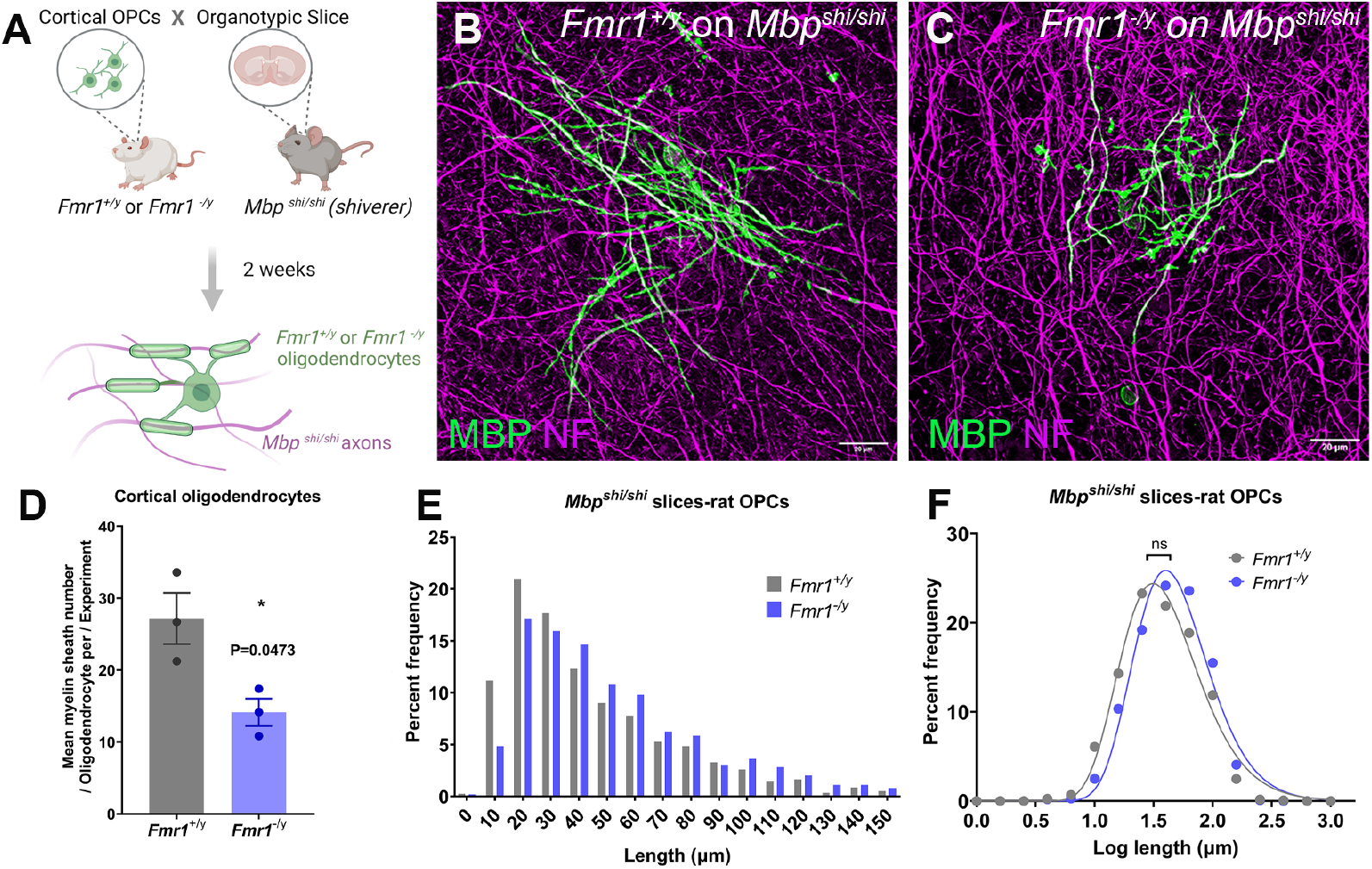
Rat *Fmr1*^*-/y*^ oligodendrocytes form fewer myelin sheaths on *Mbp*^*shi/shi*^ (Shiverer) axons *ex vivo*. **A:** Experimental design **B-C:** Representative images of *Fmr1*^*+/y*^ (B) or *Fmr1*^*-/y*^ (C) rat oligodendrocytes on mouse *Mbp*^*shi/shi*^ (shiverer) axons after 2 weeks in culture, immunostained for neurofilament (NF-magenta) and MBP (green). **D:** Mean myelin sheath number/oligodendrocyte per experiment; data presented as mean±sem and each circle is an independent experiment **E:** Histogram of sheath lengths formed by *Fmr1*^*+/y*^ or *Fmr1*^*-/y*^ rat oligodendrocytes on *Mbp*^*shi/shi*^ mouse cortical slices after 2 weeks in culture. **F:** Log sheath length plot for *Fmr1*^*+/y*^ or *Fmr1*^*-/y*^ oligodendrocytes cultured on *Mbp*^*shi/shi*^ cortical slices. Data in E and F are from 3 separate experiments per genotype. P value calculated with two-tailed, unpaired t-test with Welch’s correction in D and two-tailed unpaired t-test for the mean sheath length and mean sheath log length in E and F respectively.

These data are in accordance with the human *in vivo* transplantation experiments and together indicate that even in an *in vivo* environment with multiple *Fmr1*-expressing cell types, the maturation of *Fmr1/FMR1* null oligodendrocytes is impaired, advocating for a cell-autonomous role for *Fmr1/FMR1* in regulating oligodendrocyte maturation and morphology.

## Discussion

FXS is a common monogenic form of inherited intellectual disability and autism with extensive clinical heterogeneity (83) and cellular pathophysiology that urges the need to study the involvement of multiple cell types in the brain, as well as the inclusion of human models to improve therapeutic outcomes. Despite growing evidence of white matter abnormalities in FXS individuals (37–39) the contribution of oligodendrocyte lineage cells in FXS pathophysiology remained largely understudied.

FMRP has been previously reported to be expressed in rodent and human OPCs, in pre-myelinating and in myelinating oligodendrocytes (32, 33, 35, 36, 40, 84). We also confirmed the expression of *FMR1* mRNA in human OPCs and examined its involvement in both human and rat oligodendrocyte development. Analysis of human glial spheres showed that the generation and the proliferation of hOPCs are largely unaffected by the loss of FMRP, indicating that FMRP is not critical for hOPC production and specification. Similarly, the densities of rat oligodendrocyte lineage cells were not different between genotypes both *in vivo* and *in vitro*. A reduction in OPC cell density was previously observed in *Fmr1* knock-out mouse cerebella during the first postnatal week, while increased densities were shown in the 1-month-old mouse cortex and in the adult auditory brainstem (40, 41, 85). Furthermore, a study in zebrafish has described an increase in the number of oligodendrocyte lineage cells (SOX10+) at early larval stages in *Fmr1* knockout fish (42).

Our data indicate that although FMRP may not be critical for the generation of the oligodendrocyte lineage, it is likely important for the maturation of oligodendrocytes and the acquisition of elaborate morphologies that would allow the appropriate myelination of axons *in vitro*. Our analysis of rat and human *Fmr1/FMR1* deficient oligodendrocytes identified an impairment in oligodendrocyte maturation as indicated by the reduced MBP+/OLIG2+ cell densities in the rat mutant cultures and the significantly reduced branching networks of human and rat mutant oligodendrocytes. Oligodendrocyte morphology defects have been described to be regulated by various mechanisms (86–90), including cell-intrinsic mechanisms of the cytoskeleton and cell adhesion molecules. In fact, our bioinformatics analysis identified a variety of cytoskeletal regulators such as *Rac1, Pak1, Daam2, Sema6a*, all of which are targeted by FMRP and were previously shown to affect oligodendrocyte morphology (91–94). Given that our *in vitro* OPC cultures are devoid of axons and dendrites and several FMRP targets in the oligodendrocyte lineage are cytoskeletal proteins, it is likely that our observed morphological defects *in vitro* and possibly *in vivo* are governed by effects in the oligodendrocyte cytoskeleton.

Defects in oligodendrocyte maturation *in vivo* have also been reported in zebrafish that showed reduced *Myrf* (a marker for mature oligodendrocytes) levels in *Fmr1* null larvae (42) and in *Fmr1* knockout mice which displayed abnormal expression of myelin proteins MBP and CNPase (40). In addition, FMRP targets a number of mRNAs that have been identified as oligodendrocyte-specific regulators of myelination (e.g., synaptic proteins, cell adhesion molecules, and members of the Akt-mToR signalling pathway) that were also highly ranked in the oligodendrocyte lineage cell FMRP target expression list in our *in silico* analysis (Supplementary Figure 1, Supplementary Table 2). Synaptic-like molecular machinery is often accumulated at the points of axon-oligodendrocyte contact from the onset of myelination while oligodendrocyte-specific disruptions in this communication in both mice and in zebrafish impaired axonal myelination (59, 87, 95–98)

Although we did not observe significant differences in the densities of mature oligodendrocytes *in vivo* in rats we did observe a significant reduction in the average number of myelin sheaths they form during the third postnatal week. This defect was not accompanied by a reduction in the average myelin sheath length and was not evident at the fifth postnatal week in the rat, which suggests a myelination delay caused by the reduced formation of myelin sheaths in the medial prefrontal cortex. Nevertheless, both *Fmr1/FMR1* knockout rat and human oligodendrocytes retained this morphological impairment when transplanted onto hypomyelinated FMRP-expressing mouse axons *ex vivo* and *in vivo* respectively. This implies that cell-autonomous and conserved defects in oligodendrocytes can lead to *in vivo* delays in myelination during early postnatal development that may affect the establishment and the subsequent formation of neuronal networks. Oligodendrocytes receive inputs from neurons and respond to changes in neuronal activity through a variety of neurotransmitter receptors on their membrane (99–106). These dynamic changes are now recognised as a new form of brain plasticity that regulates the performance of neuronal networks and is important for many aspects of cognition such as social interactions (76, 96), learning and memory (77–79, 96, 107–109). Understanding how glia interact and regulate neuronal function during typical brain development is essential to reveal changes in neurodevelopmental disorders such as FXS. Our work is the first to report conserved and cell autonomous defects in both rat and FXS hPSC -derived oligodendrocytes, providing new insights into the developmental role of FMRP in human and rat oligodendrocyte maturation while offering a first glimpse into the early oligodendrocyte dysfunction in FXS.

## Acknowledgements

We would like to thank Prof Anna Williams, the members of Zoupi and Chandran groups for their comments and feedback during the writing of this manuscript. We would also like to thank the Bioresearch Veterinary Services facilities (University of Edinburgh, UK), Lynsey Dunsmore for the animal care and Dr Adrian Garcia Burgos and Dr Ian Porter from the IMPACT facility for the technical assistance. We thank the Animal Care and Resource Centre and the Central Imaging and Flow Facility at NCBS, Bangalore, India for their assistance.

## Supplementary Figures

**Fig. S1.**
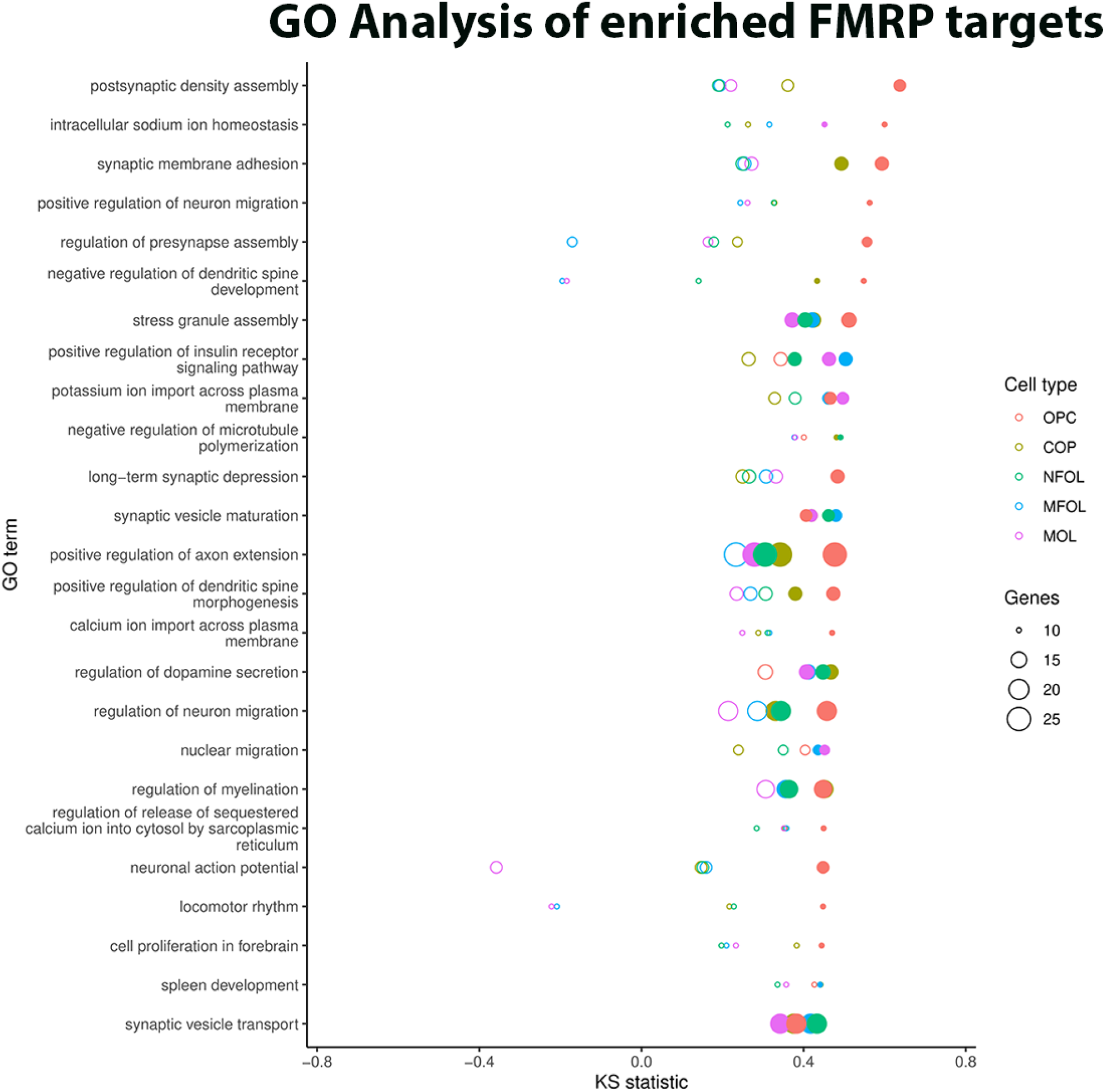
FMRP targets in rodent oligodendrocyte lineage cell classes. Graph showing the top 25 enriched GO terms from *in silico* analysis of mouse oligodendrocyte-specific datasets and FMRP gene targets in the postnatal mouse brain. GO terms were ordered by their maximum KS-test value across all cell types. Each cell type is marked with a different cell colour. The point size indicates the number of genes (i.e., FMRP mRNA targets) annotated with the GO term. Solid points indicate that the KS-test was significant at p<0.05. Oligodendrocyte clusters according to Marques et al 2016.

**Fig. S2.**
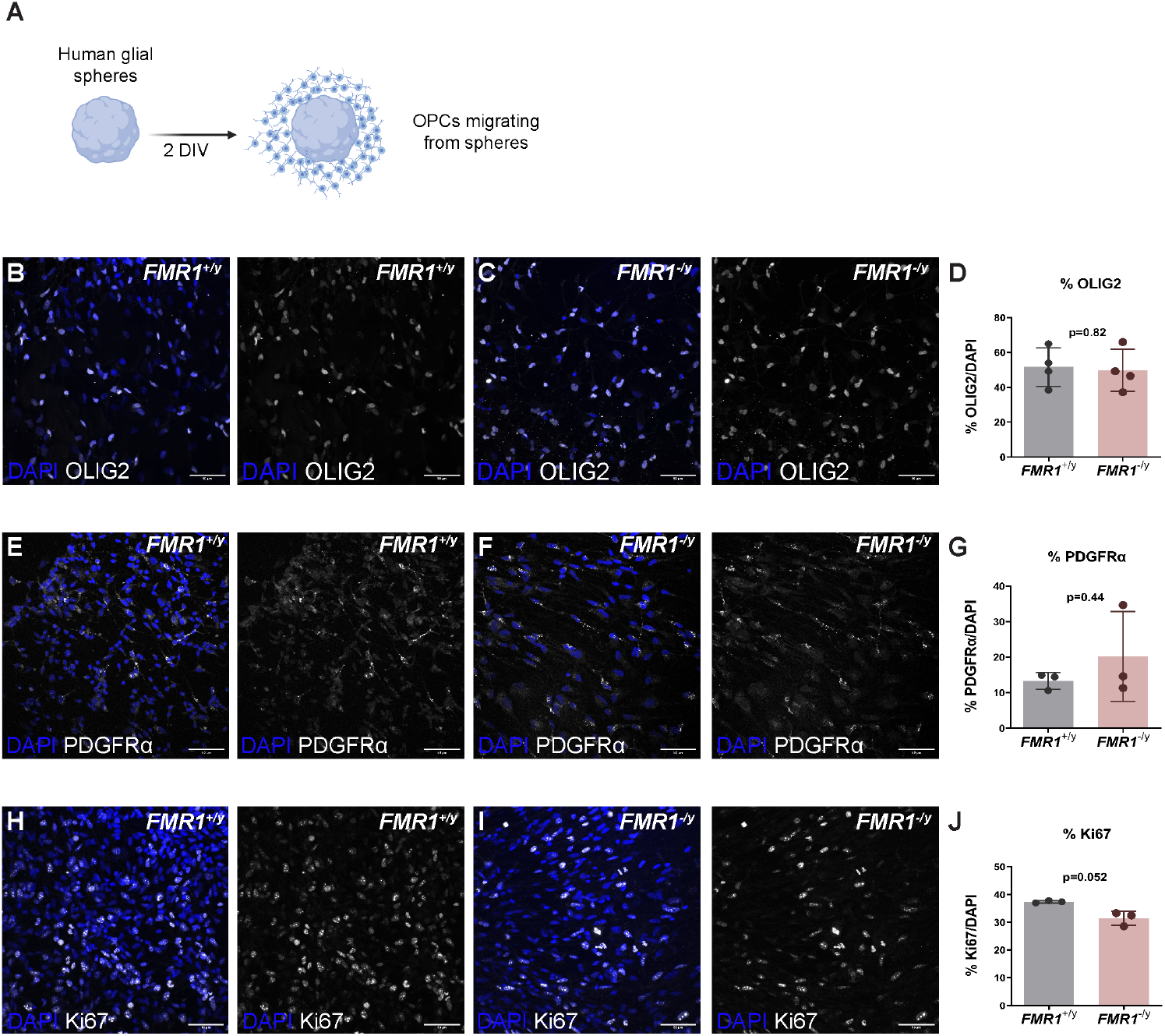
hOPC numbers and proliferation unchanged in *FMR1*^*+/y*^ and *FMR1*^*-/y*^ glial spheres. **A:** Schematic showing migration of cells from the human glial spheres. Migrating cells were enriched were OPCs and analysed in B-I panels. **B-C**,**E-F**,**H-I:** Representative images of *FMR1*^*+/y*^ and *FMR1*^*-/y*^ human oligodendrocytes after 2 days *in vitro* immunostained for OLIG2, PDGFR_α_ and Ki67 (white) and counterstained with DAPI (blue). **D**,**G**,**J:** Graph showing percent of OLIG2, PDGFR_α_ and Ki67 among total nuclei. Each data point is a different experiment. Error bars indicate sem from 3-4 different experiments; P values calculated using two-tailed unpaired t-test with Welch’s correction.

**Fig. S3.**
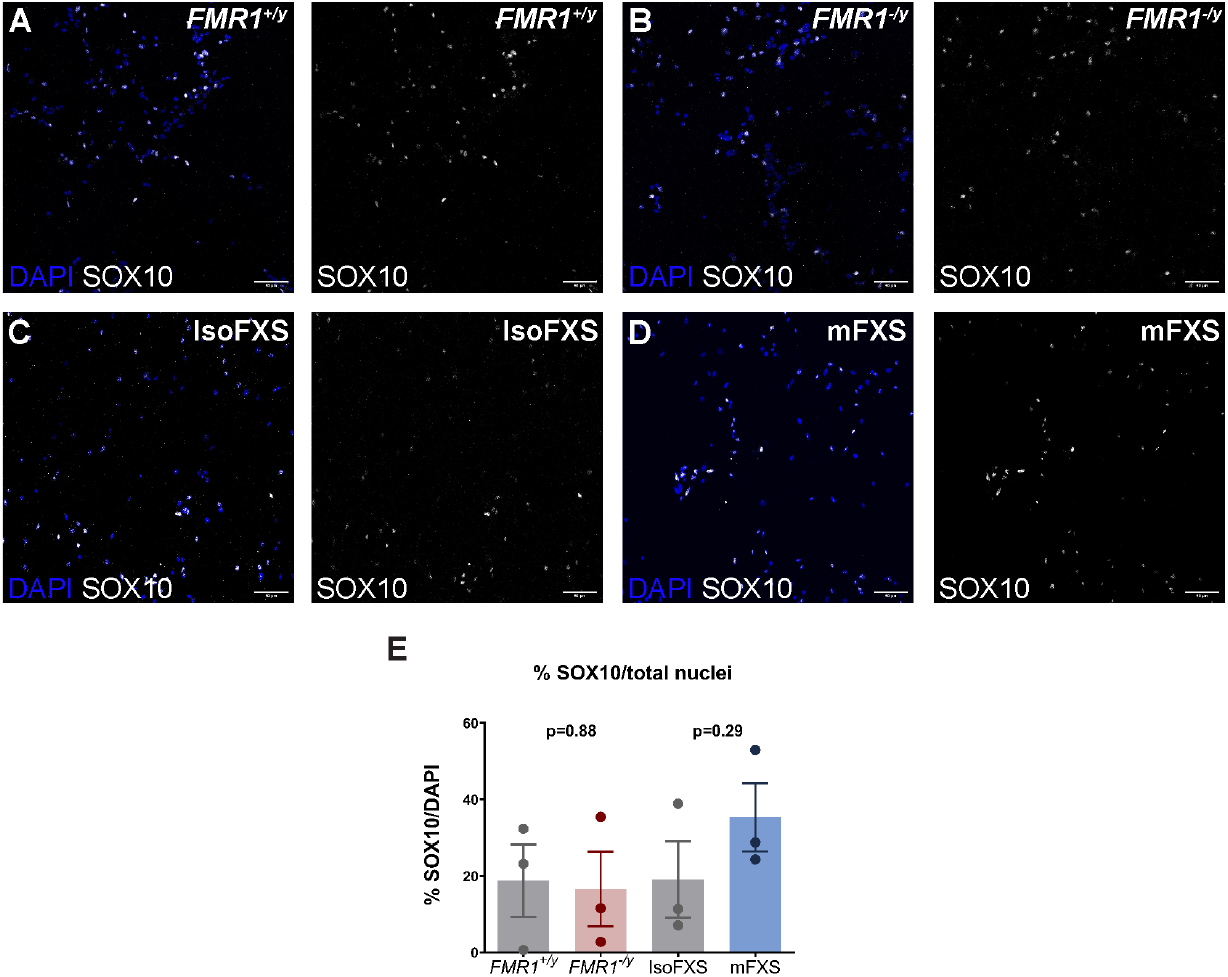
Human oligodendrocyte lineage cells unchanged in *FMR1*^*-/y*^ and mFXS glial cultures. **A-D:** Representative images of *FMR1*^*+/y*^, *FMR1*^*-/y*^, IsoFXS and mFXS Human oligodendrocytes after 7 days in vitro immunostained for SOX10 (white) and counterstained with DAPI (blue). **E:** Graph showing %SOX10 among total nuclei. Each data point is a different experiment. Error bars indicate sem from 3 different experiments; P values calculated using two-tailed unpaired t-test with Welch’s correction.

**Fig. S4.**
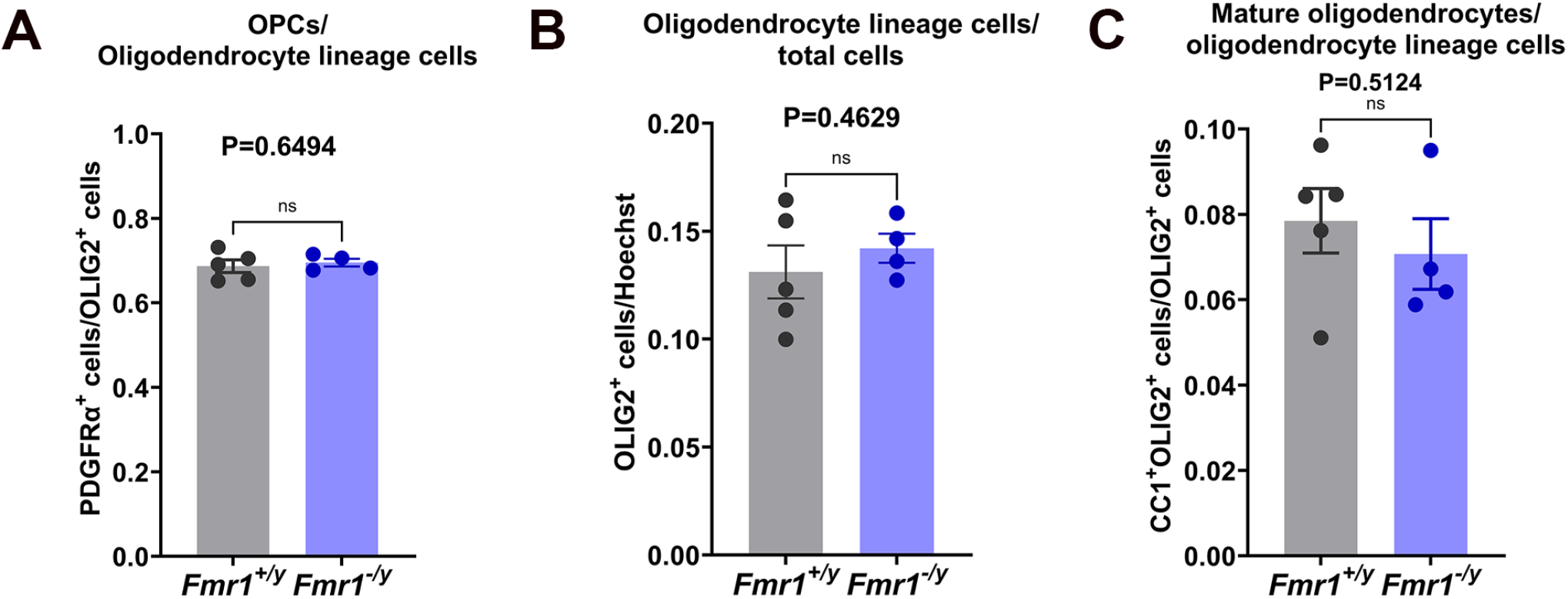
No difference in oligodendrocyte densities between genotypes in rats at the third postnatal week. **A:** Ratio of PDGFR_α_+/OLIG2+ cells over the total OLIG2+ cells in layers 2/3 of the prefrontal cortex of *Fmr1*^*+/y*^ and *Fmr1*^*-/y*^ rats at the third postnatal week **B:** Ratio of OLIG2-expressing cells over the total cell number in layers 2/3 of the prefrontal cortex of *Fmr1*^*+/y*^ and *Fmr1*^*-/y*^ rats at the third postnatal week. **C:** Ratio of CC-1+/OLIG2+ cells over the total OLIG2+ cells in layers 2/3 of the prefrontal cortex of *Fmr1*^*+/y*^ and *Fmr1*^*-/y*^ rats at the third postnatal week. Data presented as mean±sem and each circle is a rat. P values calculated with two-tailed, unpaired t-tests with Welch’s correction.

**Supplementary Table 1:**
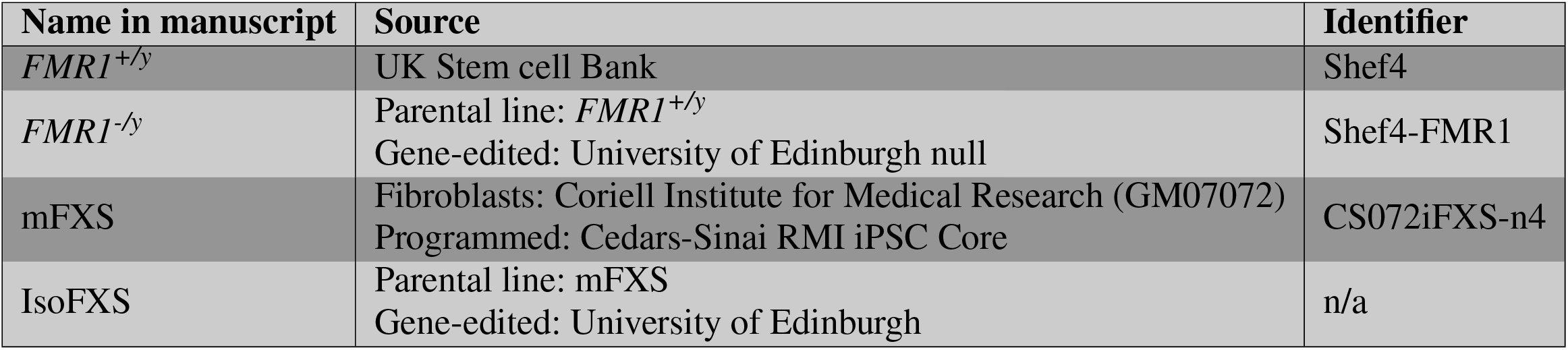
Summary of hPSC lines used in this study.

**Supplementary Table 2: excel file GO analysis data**.

